# Gestational Early-Time Restricted Feeding Results in Sex-Specific Glucose Intolerance in Adult Male Mice

**DOI:** 10.1101/2022.04.27.489576

**Authors:** Molly C. Mulcahy, Noura El Habbal, Detrick Snyder, JeAnna R. Redd, Haijing Sun, Brigid E. Gregg, Dave Bridges

**Affiliations:** University of Michigan School of Public Health, Department of Nutritional Sciences, Ann Arbor MI, USA; Michigan Medicine, Department of Pediatrics, Division of Diabetes, Endocrinology, and Metabolism, Ann Arbor MI, USA

**Keywords:** time-restricted feeding, glucose intolerance, maternal nutrition, developmental programming

## Abstract

The timing of food intake is a novel dietary component that can impact health. Time-restricted feeding (TRF), a form of intermittent fasting, manipulates food timing. During pregnancy, one may experience disruptions to food intake for diverse reasons (e.g. nausea and vomiting of pregnancy, food insecurity, desire to manage gestational weight gain, disordered eating behaviors, changes in taste and food preferences, etc) and therefore may experience periods of intentional or unintentional fasting similar to TRF protocols. Because interest in TRF is gaining popularity and feeding may be interrupted in those who are pregnant, it is important to understand the long-term effects of TRF during pregnancy on the resultant offspring. Using a mouse model, we tested the effects of gestational exposure to early TRF (eTRF) over the life course of both male and female offspring. Offspring body composition was similar between experimental groups in both males and females from weaning (day 21) to adulthood (day 70), with minor increases in food intake in eTRF females and improved glucose tolerance in males. After 10 weeks of high fat, high sucrose diet feeding, male eTRF offspring were more sensitive to insulin but developed glucose intolerance with impaired insulin secretion. As such, gestational eTRF causes sex-specific deleterious effects on glucose homeostasis after chronic high fat, high sucrose diet feeding in male offspring. Further studies are needed to determine the effect gestational eTRF has on the male pancreas as well as to elucidate the mechanisms that protect females from this metabolic dysfunction.

## Introduction

The timing of food intake in reference to circadian rhythms can impact propensity for health or disease (1). Time-restricted feeding/eating (TRF/E), a method of intermittent fasting, aligns caloric intake with naturally occurring circadian rhythms of metabolism, acting as a zeitgeber. Timing of food intake is capable of programming metabolic systems for either poor health from chronodisruption, or good health with either diurnal or nocturnal feeding, depending on the species.

To our knowledge, no estimate of the prevalence of TRE in humans exists. However, according to one sample, up to ten percent of people surveyed who state that they followed a diet in the year 2020 said they attempted “intermittent fasting,” making it the most prevalent dietary intervention in their sample (2). During pregnancy, one may have periods of time with limited food intake for many reasons: religious practice, food insecurity, disordered eating behaviors, nausea and vomiting of pregnancy/morning sickness, changes in taste/food preferences, or intentional timing of eating for weight maintenance. A recent cross-sectional study about the attitudes toward TRE in pregnant or postpartum women was conducted and found that 23.7% of those surveyed said they were willing to try TRE during pregnancy (3). The most available literature examined fasting during the month of Ramadan while pregnant. Review of these studies found that children born to those who fasted during pregnancy have babies with similar birth weights and rates of pre-term birth (4). The literature is most focused on the effects of the practice during infancy and early childhood in the resultant children.

The diet is popular and interruptions in food intake are known to occur during pregnancy; however, research about the effects of fasting during pregnancy is limited to the observance of Ramadan, a cross-sectional study about attitudes toward the practice (3), and one case report of fasting to improve gestational diabetes (5). Detailed modeling of TRF in pregnancy is warranted, as TRE exists in human populations (3, 5) and effects are unknown.

Previous studies of maternal diet during pregnancy have focused on dietary restriction or macronutrient excess in pregnancy, with little-to-no attention directed toward temporality of food intake. To date, one study of TRF during pregnancy in animals exists. This work emphasized fetal health and was completed in the context of preventing complications from overnutrition (a high fat diet, HFD) during gestation. Upadhyay and colleagues found that 9-hour TRF improved fetal lung development (6) and placental oxidative stress markers (7) at embryonic day (E)18.5 compared to ad libitum fed dams. This approach did not evaluate the long-term, postnatal effects of TRF and the independent effects of TRF are complicated by the use of a high fat diet.

The effects of TRF in non-pregnant human populations are inconsistent. Some TRF trials result in significant weight loss (8–11) while others do not (12–14). Similarly, insulin sensitization results in some (8, 14–17), but not all trials of TRF (9, 13). The way TRF is employed in human studies is rarely consistent, with varying lengths of feeding window, timing of feeding window (early vs late), control of caloric intake (isocaloric vs ad libitum feeding), inpatient observation or outpatient adherence monitoring. As such, the biological effects of this eating strategy are not clear, even in non-pregnant humans.

Results from rodent models of TRF are more consistent than human trials. These have found TRF of a HFD reduces body weight compared to ad libitum feeding (18–23), can improve Homeostatic Model Assessment for Insulin Resistance (HOMA-IR) (20, 23, 24), and may limit complications like insulin resistance (21, 22) from HFD feeding.

Taking together the likelihood that food intake can be time-disrupted in pregnancy and the evidence of TRF being a potent method to improve body composition and glycemic health in adult mice, we sought to evaluate the impact of TRF of normal laboratory chow (6-hour, early dark-cycle) before and during pregnancy on resulting offspring body composition and glycemic health through adulthood.

## Methods

### Animal care and use

Virgin female C57BL/6J mice were obtained from Jackson Laboratory (RRID IMSR_JAX:000664). All animals were maintained on a, 12-hour light/dark (12 dark (ZT12, 6pm):12 light (ZT0, 6am); ZT = zeitgeber time) cycle in a temperature and humidity-controlled room. After one week of acclimatization, they were single-housed and were assigned to feeding groups. Dams were randomized to either early time-restricted feeding (eTRF) or *ad libitum* (AL) feeding during gestation (n 8= eTRF, 9=AL). Dams fed AL had 24-hour access to a chow diet (NCD, Picolab Laboratory Rodent diet, 5L0D; 5% of Calories from fat, 24% from protein, 71% from carbohydrates). Dams fed eTRF had 6 hours of NCD food access during the early dark cycle (ZT 14-ZT 20). Water was provided *ad libitum* throughout the study to both experimental groups. After one week of either AL or eTRF feeding (beginning age 120 days), age-matched males were introduced into cages for breeding. Males were kept in the cage until a copulatory plug was detected. Daily, dams were transferred to a clean cage at ZT20, allowing for a cage free of food for eTRF animals and similar levels of handling between experimental groups. After birth, all dams switched to AL and were maintained on this diet until weaning at postnatal day (PND) 21.5. Therefore, any phenotype in the offspring is attributable to modifications to the pre-gestational/gestational diet. All experimental protocols were reviewed and approved by The University of Michigan Institutional Animal Care and Use Committee.

### Offspring growth and food intake monitoring

Pups born were weighed and counted within 24 hours of birth. Litters were reduced to 4 pups (2 male, 2 female, when possible) at PND 3.5 to standardize milk supply among litters. At PND 21.5, offspring were weighed and body composition was assessed using EchoMRI 2100 (EchoMRI) before being weaned by sex and maternal feeding regimen and housed 4-5 per cage (eTRF males = 11, eTRF females = 19, AL males = 16, AL females =17). Offspring were given AL access to NCD until PND 70. Food intake and body composition were assessed weekly. Food intake is represented as an average per animal per day. After PND 70, all animals began AL 45% High Fat Diet (HFD; Research Diets D12451; 45% Fat/ 20% Protein/ 35% Carbohydrate).

### Insulin Tolerance and Glucose Tolerance Testing

Baseline glucose (GTT) and insulin tolerance tests (ITT) were assessed at young adulthood towards the end of the NCD diet period (PND 60-70). Animals were transferred into a cage with no food during the early light cycle (ZT 2), with water freely available. After 6 hours, fasting blood glucose was assessed using a tail clip and a handheld glucometer (OneTouch Ultra). Shortly thereafter, an intraperitoneal injection of insulin was administered (Humulin, u-100; 0.75U/kg lean mass). Blood glucose was assessed by glucometer every 15 minutes for 2 hours.

One week later, glucose tolerance was assessed in a similar way (D-Glucose,1.5g/kg lean mass). Insulin and glucose tolerance were then re-assessed after HFD feeding (PND 140-160) (insulin dose 2.5U/kg lean mass, glucose dose 1.0g/kg lean mass). Area under curve was calculated for each animal by taking the sum of glucose at each time point, and then was averaged by sex and maternal feeding regimen. Rates of drop for ITT were calculated by limiting the dataset to the initial period after insulin administration (<60 minutes), taking the log of the glucose values and generating a slope for each animal. After each animal’s rate of drop was calculated, values were averaged by sex and treatment.

### Glucose Stimulated-Insulin Secretion testing in vivo

One week after GTT and ITT, animals underwent glucose stimulated insulin-secretion (GSIS) testing (PND 160-170). At ZT2, animals were placed in a clean cage without food and with unrestricted access to water. After a 6-hour fast, animals were lightly anesthetized with isoflurane via drop jar and a baseline blood sample was collected via retro-orbital bleed with a heparinized capillary. Following baseline blood collection, an intraperitoneal injection of D-glucose (1.0g/kg lean mass) was given. After 15 minutes, animals were lightly anesthetized in the same manner and another blood sample was collected. Blood samples were allowed to clot on wet ice (∼20 minutes), then were spun down in a cold centrifuge (4 °C, Eppendorf microcentrifuge, model 5415R) for 20 minutes at 2000 g. Serum was pipetted off and stored at - 80 °C until analysis. Serum insulin was assessed via a commercially available ELISA kit (ALPCO 80-INSMSU-E10). Insulin was assessed in 5uL of serum and read via colorimetric assay.

### Statistical analysis

All measures with p-values <0.05 were considered statistically significant. Data are presented as mean +/- standard error throughout. All statistical analyses were performed using R version 4.0.2 (25). Repeated measures, such as body composition, cumulative food intake, and responses to GTT or ITT were assessed via mixed linear effects modeling with random effects of mouse ID and dam and fixed effects of maternal dietary treatment, age, and sex using lme4 version 1.1-26 (26). Body composition and food intake were measured separately in 2 phases: during NCD feeding, and after being switched to HFD. Analyses were tested for significant interactions between sex and maternal dietary treatment. Models were assessed using a two-way ANOVA for sex and maternal dietary treatment, with an interaction between the two. If a significant interaction was observed, sex-stratified models were then used and the p-value for the interaction was reported. Otherwise, sex was used as a covariate in a non-interacting model. Observations were tested for normality by Shapiro-Wilk test and equivalence of variance by Levene’s test. Pairwise measures that were normal and of equal variance utilized Student’s *t*- tests. Measures that were not normally distributed used non-parametric Mann-Whitney tests.

## Results

### Gestational eTRF increases food intake, but not body composition in early life

To model gestational early time restricted feeding (eTRF), we used a normal chow diet (NCD) and assigned female mice to either unrestricted (*ad libitum*, AL) or 6 hours of restricted food availability between ZT14-20 (eTRF, 50% of their active nocturnal window) (**Figure 1A**). This approach limits potential sleep disruptions and is more translationally relevant to human dietary restriction. This treatment started a week before mating and continued through delivery (**Figure 1B**). Litters were normalized to equal sizes to reduce variability and effects of lactation.

**Figure 1:**
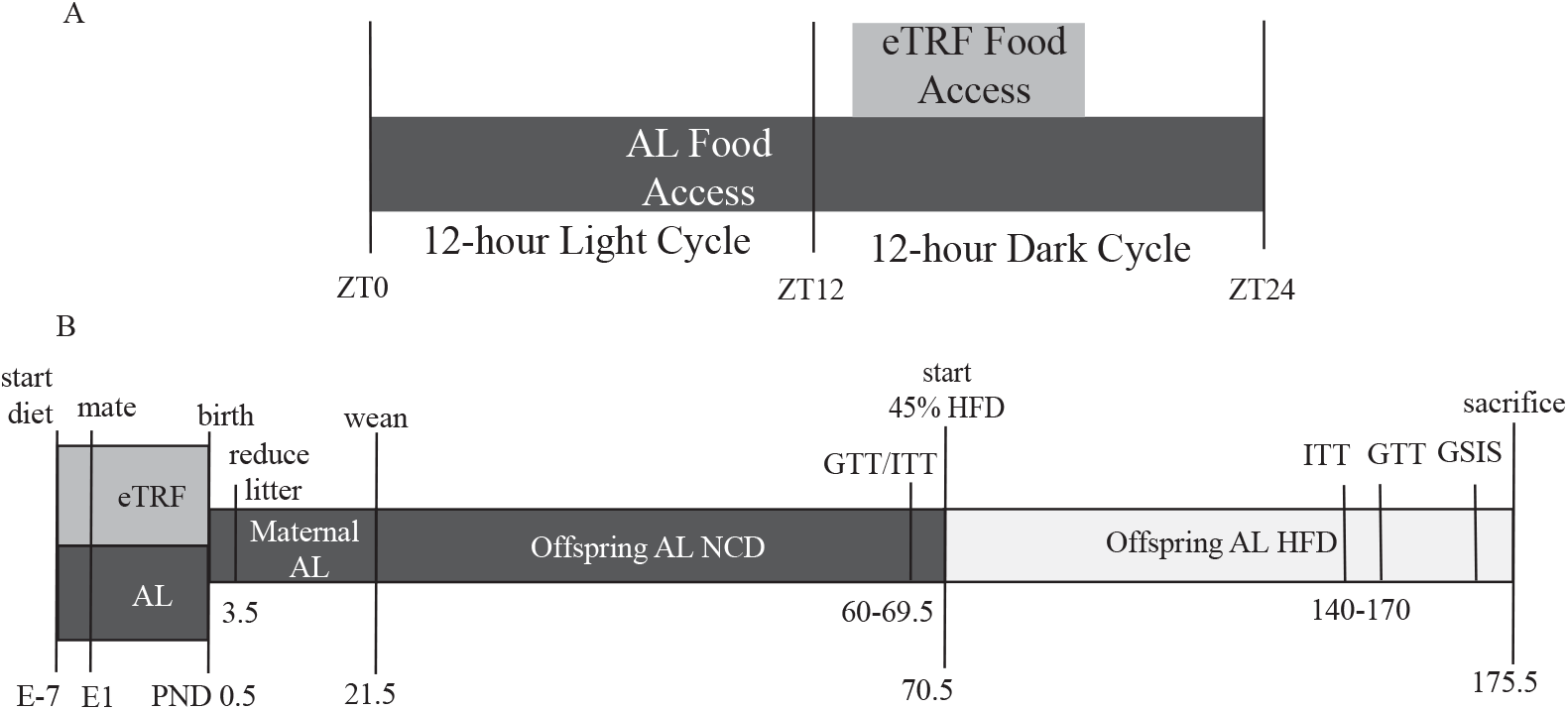
Experimental Protocol and Timing. A) Food availability and timing for dams during pregnancy. Food access began at ZT13 for early Time-Restricted Feeding dams (eTRF, light gray, n=8) and continued until ZT129, total of 6 hours. Food was available 24 hours a day for ad libitum dams (AL, dark gray, n=9). B) Offspring experimental protocol. After birth, all dams had AL access to laboratory chow (NCD). Litters were reduced to 4 (2 males, 2 females when possible) on post-natal day (PND) 3. Offspring were weaned by maternal feeding regimen at PND 21 and maintained on AL NCD for 70 days. Weekly body composition and food intake measurements were taken throughout the experiment. At 70 days of age, insulin tolerance tests (ITT) and glucose tolerance tests (GTT) were conducted before switching all animals to a 45% high fat diet (HFD) with sucrose. Animals were on HFD for 10 weeks before repeating ITT and GTT, and an in vivo glucose stimulated insulin secretion test (GSIS). Animals were euthanized after these tests. Abbreviations: zeitgeber time (ZT), ZT0 = lights on, ZT12 = lights off.

The pups were weighed and their body composition was assessed weekly, then analyzed using linear mixed effect modeling. We found significant and expected effects of age and sex (older mice weigh more than younger mice and male pups weigh more than females), but no effect modification of maternal eTRF on body weight (**Figure 2A**, p_diet_=0.47), lean mass (**Figure 2C**, p_diet_=0.45), or fat mass (**Figure 2B**, p_diet_=0.47). There was no interaction between sex and maternal intervention in cumulative food intake (p_diet*sex_=0.38). However, cumulative food intake in the NCD period is 22% higher in eTRF females than AL females and 10% higher in eTRF males than AL males (**Figure 2D**, p_diet =_ 0.016). Assessing the efficiency by which food is converted into stored mass resulted in a 12% reduced feeding efficiency in eTRF female offspring (p_sex_<0.00001) which is not present in males (**Supplementary Figure 1A**).

**Figure 2:**
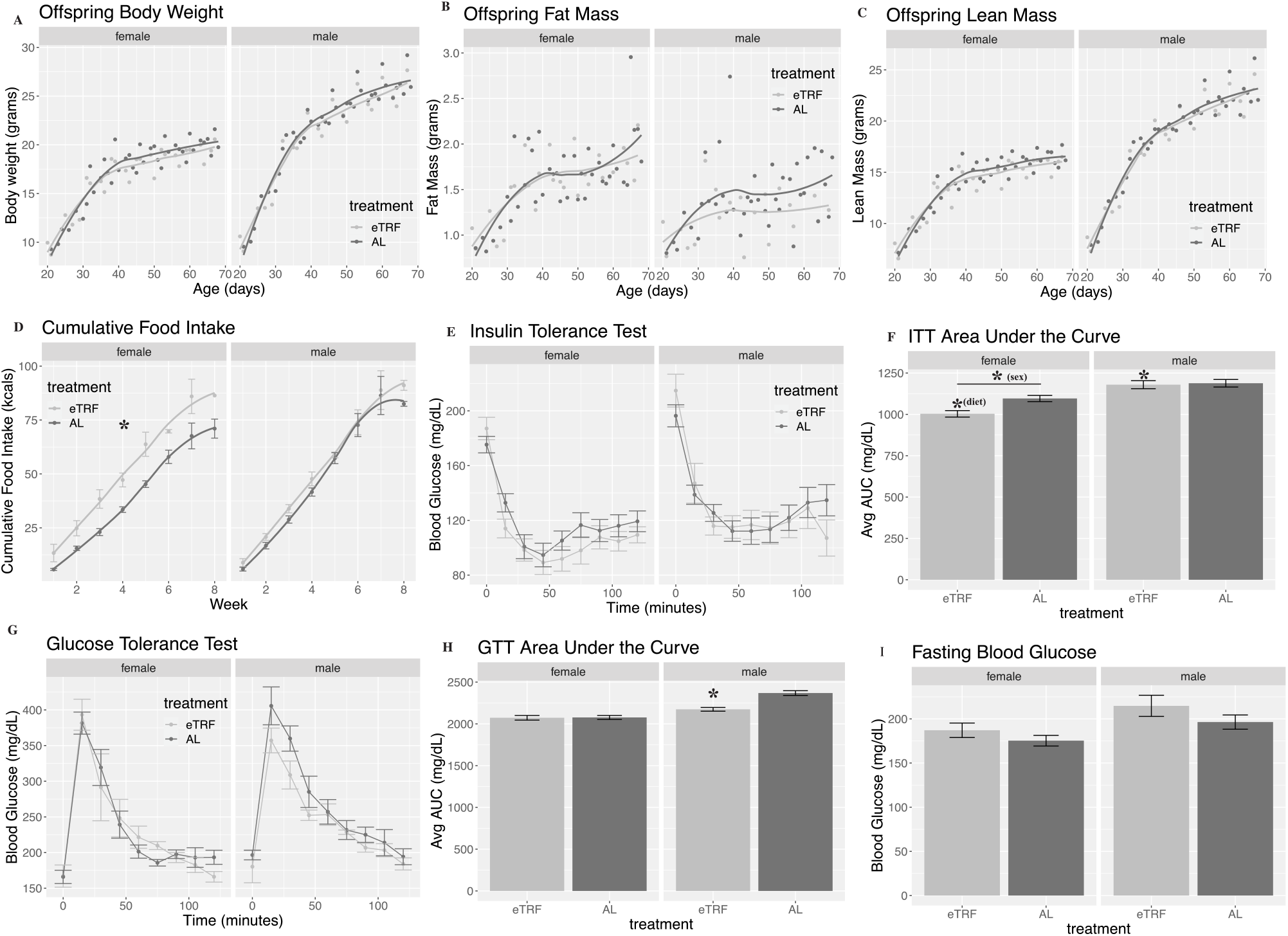
Early Life Body Composition, Food Intake, and Glycemic Homeostasis. **A)** Body weight in grams from PND21-PND70 in males and females, averaged by age, maternal feeding regimen, and sex. **B)** Fat mass in grams from PND21-PND70 in males and females, averaged by age, maternal feeding regimen, and sex. **C)** Lean mass in grams from PND21-PND70 in males and females, averaged by age, maternal feeding regimen, and sex. **D)** Food intake in kcals per mouse per day, averaged by week, maternal feeding regimen, and sex. *p-value <0.05 for diet. **E)** Insulin tolerance test (ITT) ∼PND 70, averaged by maternal feeding regimen, sex, and time in minutes. **F)** Area under the curve (AUC) for ITT, averaged by maternal feeding regimen, and sex. * indicates p-value <0.05 for effect of diet in males. **G)** Glucose tolerance test (GTT) ∼PNG 70, averaged by maternal feeding regimen, sex, and time in minutes. **H)** AUC for GTT, averaged by maternal feeding regimen, and sex. * indicates p-value <0.05 for effect of diet in males. **I)** Fasting blood glucose (FBG) PND 70, averaged by maternal feeding regimen and sex. Animals included in body composition measurements, FBG, ITT, and GTT, n=11 eTRF males, 16 AL males, 19 eTRF females, 17 AL females. Number of cages in food intake analysis n=4 eTRF males, 5 AL males, 4 eTRF females, 5 AL females.

### Gestational eTRF modestly improves glucose tolerance in young adult males

To assess glucose homeostasis in the offspring, we conducted ITTs and GTTs between PND 60 and 70. Male offspring averaged 15mg/dL higher blood glucose during insulin tolerance testing compared to females (p_sex_=0.0018), but no effect of maternal dietary restriction was evident through linear mixed effect modeling (**Figure 2E**, p_diet_=0.73). Summarizing the ITT by calculating the area under the curve (AUC) demonstrated there was no diet:sex interaction (p_diet:sex_=0.069), but an effect of maternal restriction where eTRF offspring had lower AUC compared to AL offspring, 8.5% and 2.2% lower in females and males respectively (p_diet_=0.013), and a significant effect of sex (p_sex_<0.0001). As expected, males had a higher AUC than females (**Figure 2F**). The initial response to insulin (the rate of glucose decline over the first 60 minutes, not pictured) was not significant for sex (p_sex_=0.10) or treatment (p_diet_=0.83). Taken together these data suggest that gestational eTRF slightly improves the response to insulin challenge in adult mice, and that this is not driven by increased fat mass.

Glucose tolerance was similar in young adulthood between groups in both males and females (**Figure 2G**). We found no significant effect of diet (p_diet_=0.53) on the rise in blood glucose during GTT, but there was an effect of sex (p_sex_=0.0093) on glucose tolerance, again with expected higher glucose levels in male mice. The summarized AUC for the GTT (**Figure 2H**) shows a significant interaction between sex and maternal dietary treatment (p_sex:diet_=0.00082). eTRF males had an 8.2% lower AUC than their AL counterparts (p_diet_<0.0001) while this was absent in females (p_diet_=0.99). Fasting blood glucose, assessed before ITT and GTT, was 10.4% higher in males than in females (p_sex_=0.0054; **Figure 2I**), but did not differ significantly by maternal dietary treatment (p_diet_=0.18). Thus, there are modest glycemic effects of gestational eTRF present in young, chow-fed male offspring that were not explained by differences in weight or body composition, which was comparable between groups.

### HFD feeding in adult offspring exposed to eTRF during gestation generates sex-specific glucose intolerance

Given that adult offspring were minimally affected by gestational eTRF exposure, we administered a high fat, high sucrose (HFD) overnutrition challenge; *ad libitum* access to 45% of energy from fat and 17% of energy from sucrose after PND 70. Food intake and body composition measurements continued weekly. Similar to the findings on chow, with HFD, there were no major differences between eTRF and AL offspring in body weight (**Figure 3A**, p_diet_=0.99), fat mass (**Figure 3B**, p_diet_=0.65), or lean mass (**Figure 3C**, p_diet_=0.47). Therefore, offspring of eTRF and AL experienced similar changes in body composition in response to overnutrition. Cumulative HFD consumption was comparable between females and males (p_sex_=0.72), and maternal restriction groups (**Figure 3D**, p_diet_=0.72). Feeding efficiency was greater in males than in females, which is consistent with the NCD period (**Supplemental Figure 1B**, p_sex_ = 0.00023). However, unlike the NCD period, efficiency was indistinguishable between eTRF and AL offspring (p_diet_=0.93).

**Figure 3:**
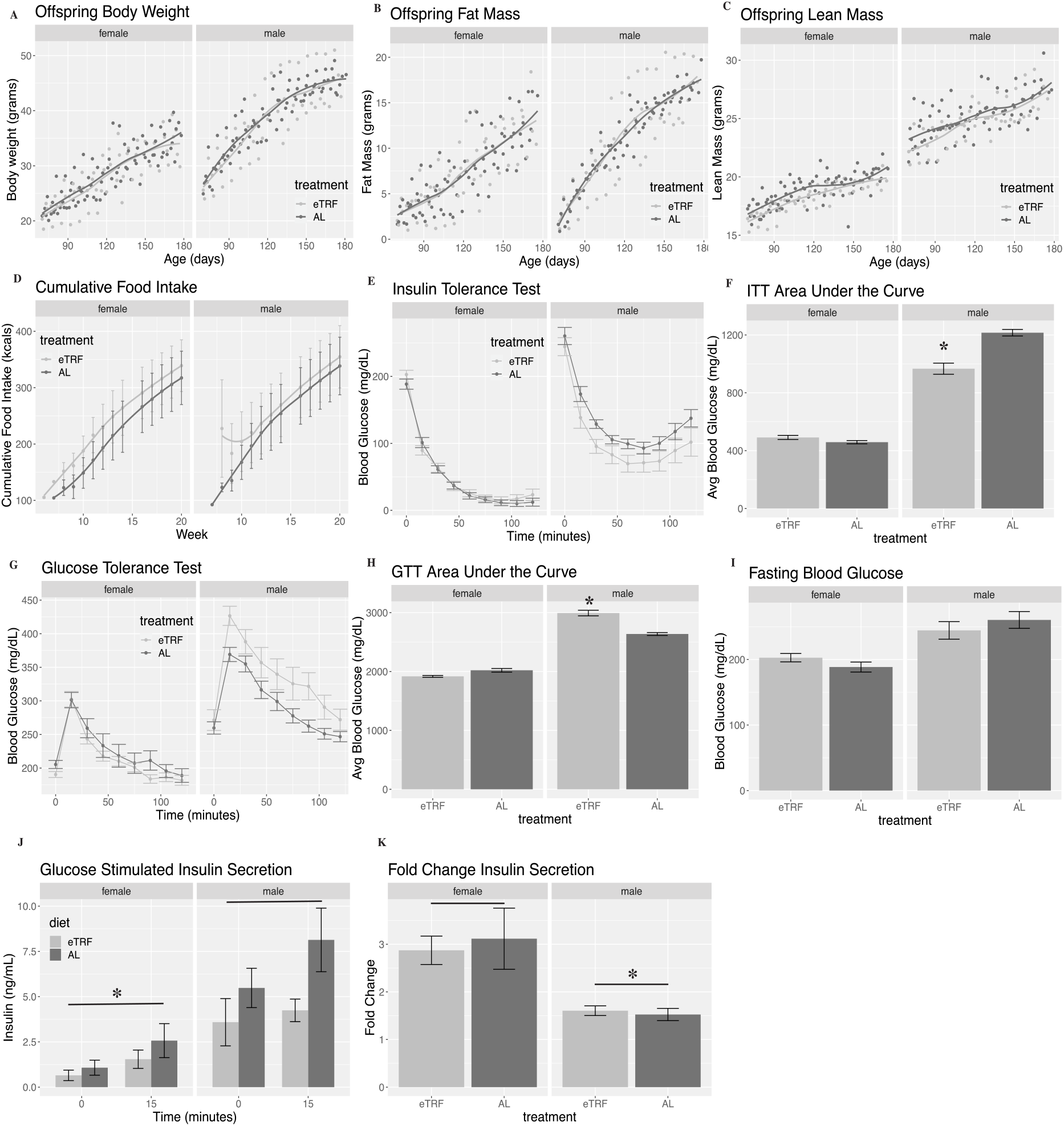
Body Composition, Food Intake, and Glycemic Response to High Fat Diet Feeding in Adulthood. **A)** Body weight in grams from PND 70-175 in males and females, averaged by age, maternal feeding regimen, and sex. **B)** Fat mass in grams from PND 70-175 in males and females, averaged by age, maternal feeding regimen, and sex. **C)** Lean mass in grams from PND 70-175 in males and females, averaged by age, maternal feeding regimen, and sex. **D)** High fat diet (HFD) intake in kcals per mouse per day averaged by week, maternal feeding regimen, and sex. **E)** Insulin tolerance test (ITT) after 10 week of HFD, averaged by age, maternal feeding regimen, sex, and time in minutes. **F)** Area under the curve (AUC) for insulin tolerance test, averaged by maternal feeding regimen, and sex. * indicates, p-value <0.05 for diet in males. **G)** Glucose tolerance test (GTT) after 10 weeks of HFD, averaged by maternal feeding regimen, sex and time in minutes. **H)** Area under the curve (AUC) for GTT after 10 weeks of HFD, averaged by maternal feeding regimen and sex. * indicates p-value <0.05 for effect of diet in males. **I)** Fasting blood glucose (FBG) after 10 weeks HFD, averaged by maternal feeding regimen, and sex. **J)** Glucose stimulated insulin secretion (GSIS), averaged by maternal feeding regiment, sex, and time. * indicates p-value <0.05 for effect of sex. Animals included in body composition, FBG, ITT, GTT, and GSIS: n=11 eTRF males, 16 AL males, 19 eTRF females, 17 AL females. Cages in food intake analysis: n=4 eTRF males, 5 AL males, 4 eTRF females, 5 AL females.

We repeated an ITT and GTT after 10 weeks of HFD feeding. During the ITT, there was a significant interaction between sex and diet using mixed linear effect modeling (**Figure 3E**, p_sex:diet_=0.03). Female eTRF had a similar response to insulin, with less than a 1 mg/dL difference from their AL counterparts (p_diet_=0.85), but male eTRF offspring tended to be more insulin sensitive with 25mg/dL lower glucose compared to AL males (p_diet_=0.17). These findings were confirmed by calculating the AUC where eTRF females had 7% greater AUC than AL females (**Figure 3F**, p_diet_=0.20) while eTRF males had 20.4% lower AUC than AL males (p_diet_<0.0001). The initial rate of glucose decline (not pictured) was greater in females compared to males (p_sex_=0.029) but there were no differences between eTRF and AL offspring (p_diet_=0.23). The trend toward insulin sensitivity from the ITT was not explained by fasting blood glucose, as females had 23% lower fasting blood glucose than males (p_sex_<0.0001) but were similar between eTRF and AL offspring within the same sex (**Figure 3I**, p_diet_=0.83). Glucose tolerance tests in **Figure 3G**, also showed significant effect of interaction (p_sex:diet_=0.011), although now in the opposite direction. During GTT, eTRF males trended toward glucose intolerance with an average of 53mg/dL higher glucose than AL males during the course of the experiment (p_diet_=0.14). This was not observed in female eTRF offspring, which had similar blood glucose during the GTT compared to AL females (p_diet_=0.61). The GTT AUC showed interaction between effects of sex and treatment (**Figure 3H**, (p_sex:diet_<0.0001)). AUC was 5% lower in eTRF females (p_diet_=0.07) but was 13.5% higher in eTRF male offspring compared to AL (p_diet_<0.0001). Taken together, these tests suggest eTRF causes male-specific glucose intolerance and insulin sensitivity. Given that we cannot explain glucose intolerance in males via reduced insulin sensitivity, we next evaluated insulin secretion.

To test for insulin secretion defects, we conducted an *in vivo* glucose stimulated insulin secretion (GSIS) assay (**Figure 3J**). Females had lower levels of insulin than males (p_sex_<0.0001). There was a non-significant trend toward lower insulin levels in eTRF compared to AL offspring of both sexes (p_diet_=0.071). Females had similar increases in insulin in response to glucose injection, 139% in AL versus 137% eTRF. Male AL offspring had a 48% increase in insulin whereas this was just an 18% increase for eTRF males. There was no interaction between sex and maternal restriction (p_sex:diet_=0.064). Females have 94% greater fold-change insulin secretion in response to glucose challenge than male offspring (p_sex_=0.0027) and there is no impact of maternal restriction (p=0.85, **Figure 3K**). Although not conclusive, the GSIS lends support to a model that sex-specific defects in insulin secretion result in sex-specific glucose intolerance after HFD challenge in males exposed to eTRF *in utero*.

## Discussion

This study is the first to describe the long-term effects of gestational eTRF on offspring health and their response to a high fat, high sucrose diet challenge. We find significant deleterious effects of gestational eTRF on glucose tolerance are present only in adult male offspring when exposed to long-term HFD feeding. Based on GSIS testing, we propose that this is attributable to impaired insulin secretion, as insulin secretion tended to be lower in eTRF males compared to their AL counterparts, although this did not reach statistical significance. Other studies of TRF using HFD in mice provide evidence that fasting insulin is lowered (20–23, 27) and resulting HOMA-IR is improved (22, 27, 28). We see that baseline insulin is modestly lower in male offspring, and this could contribute to the modest insulin sensitivity seen after HFD feeding. Our finding that fasting blood glucose is unchanged in eTRF compared to AL exposed mice is confirmed by other groups examining TRF with HFD (20, 27, 28). The elevated food intake in female offspring exposed to eTRF *in utero* is novel. Studies of adult mice pairing TRF and HFD report reduced food intake (24, 29) or equivalent caloric intake when matched by diet (21–23, 30). This could indicate a compensatory response in the female offspring resulting from eTRF *in utero*. Interestingly, this did not result in differing body weight or composition, suggesting that this increased food intake is matched by decreased caloric extraction or increased energy expenditure in these mice.

The phenotype in male offspring from this study bears resemblance to animal models of intrauterine growth restriction (IUGR), where glucose intolerance in resultant offspring is common. First described by Barker and colleagues, offspring who were deprived of nutrition *in utero* were more likely to develop chronic, nutrition-related disease in adulthood (31). Since that time, multiple animal models for IUGR were developed; maternal overnutrition during pregnancy, maternal caloric restriction, maternal protein restriction, and surgically induced placental insufficiency through late gestation uterine artery ligation. Undernutrition in pregnancy often results in offspring development of glucose intolerance (32–34). The extent to which male-specific effects are seen is difficult to deduce as many groups either study male offspring exclusively (34, 35) or analyze males and females together (33, 36). Male offspring who had placental insufficiency develop glucose intolerance in adulthood (37, 38), but females can also develop glucose intolerance (39, 40). Maternal overnutrition can also result in males with glucose intolerance (41, 42). Therefore, metabolic effects being limited in the current study to male offspring is consistent with much of the literature, as females appear to be less affected.

Glucose intolerance in IUGR models often occurs with insulin-related abnormalities in the offspring, such as lower insulin content in the pancreas (33), lower basal insulin levels (36), impaired insulin secretion (34, 40), increased pancreatic islet size (42), altered vascularity of islets (43), or reduced beta cell mass (44). These abnormalities are also accompanied by abnormal glucose tolerance in adulthood (32, 42). However, in the present study we find modest improvement in male insulin sensitivity in adulthood in male offspring exposed to gestational eTRF. We attribute male-specific insulin sensitivity during high fat diet feeding to eTRF males having lower basal levels of insulin compared to AL males. This means that peripheral tissues would be more sensitive to insulin action despite an apparent insulin secretion impairment at the level of the pancreas. The similarity of the present study to those using diverse gestational stressors suggests that restriction of the total time of food intake in dams is sufficient to induce offspring glucose intolerance similar to IUGR models, but not insulin resistance.

Although we have not investigated offspring pancreatic tissues, we hypothesize that alterations in the development of the pancreas may underlie the male-specific glucose intolerance and modest insulin sensitivity in eTRF offspring. Time-limiting the availability of nutrients to the fetus through eTRF may program offspring pancreas for development during daily periods of nutrient scarcity and result in impaired beta cell development or islet size leading to reduced insulin secretion. Intrinsic changes in islet function are also possible. Studies done in adult male animals undergoing TRF with chronodisruption have also found that time-restricting food access reduced insulin production with secretion most affected (enhanced compared to controls) and found no effect of insulin tolerance (45). This is confirmed by one study of early post-natal exposure to TRF, which found that adolescent males who were fed TRF the first 4 weeks after weaning developed smaller islets of Langerhans and higher blood glucose compared to those fed AL (30). Another contributor to this phenomenon may be that the islets were able to compensate in young male offspring during a lower-calorie diet (NCD) and therefore the effect did not become apparent until an overnutrition challenge during adulthood. Therefore, future studies of gestational or developmental eTRF should examine islet size, pancreatic beta cell mass, and insulin secretion to investigate the mechanism of offspring glucose intolerance further.

This study and the conclusions to be made from it have some limitations. First, the model of gestational eTRF may have resulted in differences in maternal behaviors that were not noted by the study team, and therefore could play a part in the effects seen in the offspring. Second, although we see a robust effect on glucose intolerance and trends of lower insulin secretion in male eTRF offspring in adulthood, we did not evaluate islet size or beta cell mass to determine the mechanism driving the worsening of glucose tolerance in adulthood.

There are many strengths to this study. Among them are the use of a preclinical model which facilitates consistency when compared to existing literature. Further strengths include the long follow up period for a gestational exposure, controlling for the effect of litter size, repeated measurement of body composition, and food intake measurements over the life course in the resultant offspring. Finally, the inclusion of both male and female offspring in the study, as many metabolic assessments of TRF either focus exclusively on the effects of the regimen in males (22, 23) or female mice (20, 21) is a strength.

Future work with this model should include assessment of offspring pancreatic tissue. Finally, our model used healthy non-obese dams and our results cannot be extended to effects of eTRF in the context of metabolic syndrome, diabetes, or obesity during pregnancy.

## Conclusion

Offspring who are exposed to eTRF of NCD *in utero* have similar body composition, glucose tolerance, and insulin tolerance in early adulthood in both males and females. Gestational eTRF led to sex-specific impairments in male glucose tolerance in adulthood after chronic HFD feeding, likely due to impaired insulin secretion. This occurs without increase in body weight, fat mass, or food intake compared to age matched AL males. More research is warranted to understand the mechanisms that underlie this novel phenotype.

---

**Supplemental Figure 1:**
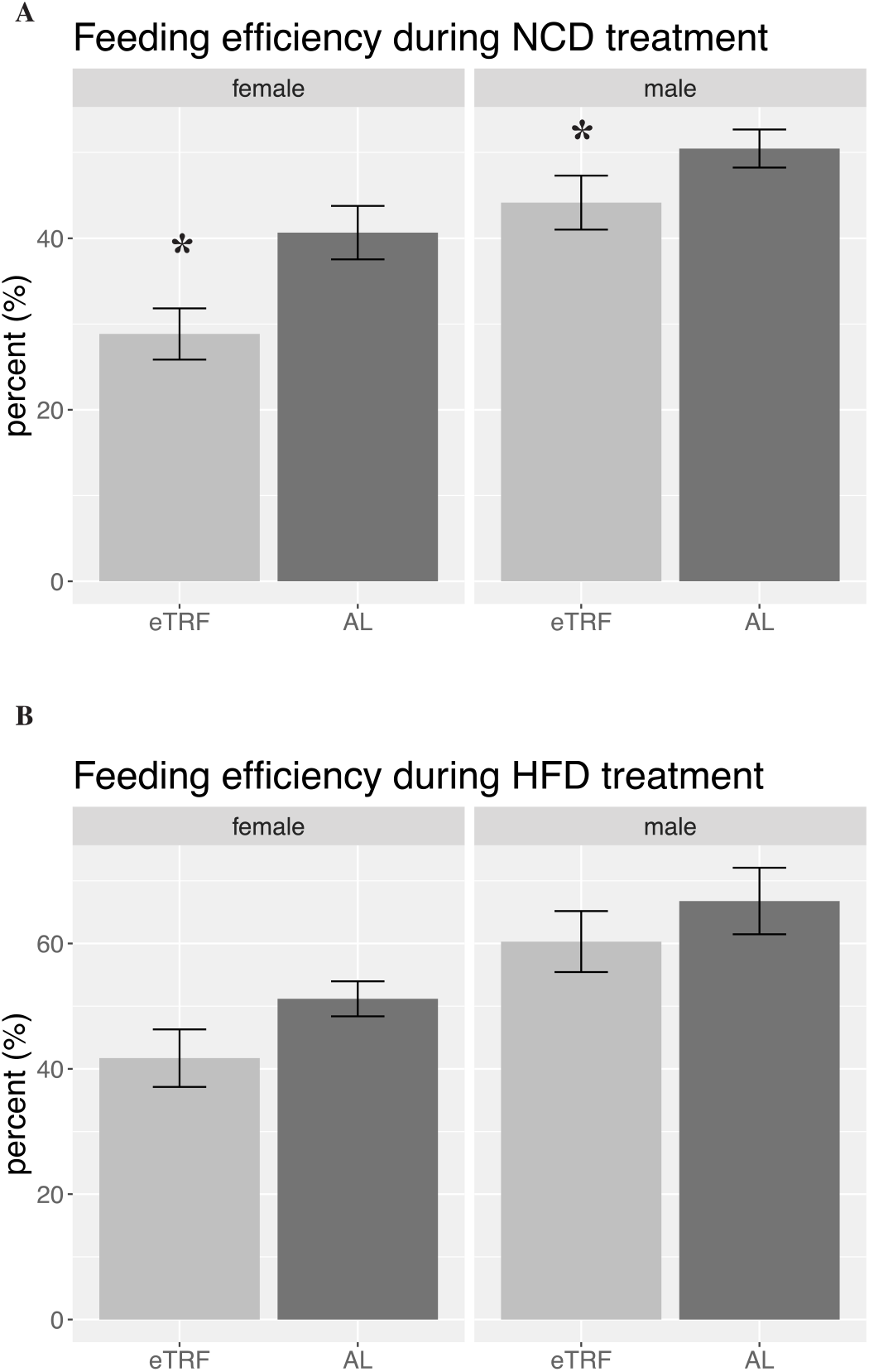
Feeding Efficiency Throughout Adulthood. **A)** Feeding efficiency (%) in males and females, calculated based on food intake and body composition changes during the NCD period (before PND 70). (p_sex_<0.001, p_diet_=0.002). **B)** Feeding efficiency in males and females during the HFD period (after PND 70). (p_sex_ = 0.00023, p_diet_ = 0.093).

